# Pair bond endurance promotes cooperative food defense and inhibits conflict in coral reef butterflyfishes

**DOI:** 10.1101/214627

**Authors:** Jessica P. Nowicki, Stefan P. W. Walker, Darren J. Coker, Andrew S. Hoey, Katia J. Nicolet, Morgan S. Pratchett

## Abstract

Pair bonding is generally linked to monogamous mating systems, where the reproductive benefits of extended mate guarding and/or of bi-parental care are considered key adaptive functions. However, in some species, including coral reef butterflyfishes (f. Chaetodonitidae), pair bonding occurs in sexually immature and homosexual partners, and in the absence of parental care, suggesting there must be non-reproductive adaptive benefits of pair bonding. Here, we examined whether pair bonding butterflyfishes cooperate in defense of food, conferring direct benefits to one or both partners. Pairs of *Chaetodon lunulatus* and *C. baronessa* use contrasting cooperative strategies. In *C. lunulatus,* both partners mutually defend their territory, while in *C. baronessa,* males prioritize territory defence; conferring improvements in feeding and energy reserves in both sexes relative to solitary counterparts. We further demonstrate that partner fidelity contributes to this function by showing that re-pairing invokes intra-pair conflict and inhibits cooperatively-derived feeding benefits, and that partner endurance is required for these costs to abate. Overall, our results suggest that in butterflyfishes, pair bonding enhances cooperative defense of prey resources, ultimately benefiting both partners by improving food resource acquisition and energy reserves.

## Introduction

Pair bonding, a selective pro-social and enduring affiliation between two individuals that is maintained beyond reproduction, has independently evolved numerous times across the animal kingdom^1-3^. Pair bonding is generally associated with monogamous mating (mammals^4^, birds^5^, reptiles^6^, amphibians^7^, marine fishes^3^) where it has been hypothesised/shown to be advantageous due to reproductive benefits of extended mate-guarding^8,9^ and/or bi-parental care^10^. However, the presence of pair bonding between sexually immature^11^ and homosexual^12,13^ partners indicates that the benefits of pairing extend beyond those of reproduction.

Aside from mate-guarding and bi-parental care, pair bonding might be attributed to the benefits of social assistance during ecological processes that are directly conferred to one or both partners^14–16^. One such process may be cooperative defense of high value resources; such as food, shelter, or nesting sites^17,18^; by one or both partners. In heterosexual pairs, resources are often defended primarily or exclusively by males (*sensu* male-prioritized “division of labor”^19^ or “resource brokering”^20^), with benefits presumably related to increased mating access to^21^ or fecundity of^22^ females, or to other resources/services that are partitioned by females (e.g., burrow maintenance)^19,23^. Alternatively, resources may be mutually defended, or “co-defended” by male and female partners^11,21^, presumably because both partners directly benefit from sharing this responsibility^21^.

This cooperative or assisted resource defense hypothesis (ARDH) for pair bonding makes several fundamental predictions. Male-prioritized defense expects that males primarily defend resources within a territory^3,17,21^ wherein females are unable to maintain a territory alone and/or directly benefit from male’s assistance^3^. Alternatively, mutual resource defense predicts that both partners mutually defend resources within a territory^3,17,21,23^, wherein both are unable to maintain a territory alone and/or directly benefit from each other’s assistance^3,23^. Although the role of assisted resource defense (ARD) in promoting pair bonding has received less research attention than that of mate-guarding or bi-parental care, *in situ* observations and explicit tests of these predictions have supported the ARDH for pair bonding across a wide range of taxa (Supplementary Table S1 online) (but see^24,25^).

Butterflyfishes of the genus *Chaetodon* are ideal model taxa for testing the ARDH for pair bonding. Among the ~ 91 species within the genus, at least 59 reportedly pair bond (data sourced from^11,26–29^. Heterosexual pairs of at least some species (mainly, *Chaetodon lunulatus*) are monogamous^30,31^ and display mate-guarding^11,32^. However, same-sexed^12,33^, reproductively immature^15,33^ and reproductively inactive^11,34^ pairing also occurs. Moreover, butterflyfishes do not provide parental care^30,35,36^. At least 24 *Chaetodon* species feed predominantly (≥ 80%), if not exclusively, on a diet that is temporally and spatially stable, and therefore economically defendable (i.e., coral)^32,37,38^. There is, however, considerable variation in the level of dietary specialization among corallivorous butterflyfishes that is related to interspecific dominance over feeding sites, such that obligate corallivores dominate territorial disputes over feeding generalists^39^. Pairing in corallivorous butterflyfishes is suggested to arise from the need for assisted defense of coral prey against con-and heterospecifics in order to better invest in feeding and energy reserves^11,40,41^. Although pair bonding butterflyfishes are presumed to have very high levels of partner fidelity (up to 7 yrs) (Supplementary Table S2 online), the ecological basis of pair bond fidelity among these organisms remains unknown.

The overall aim of this study was to test whether pair bonding in two species of common coral-feeding butterflyfishes (*C. lunulatus* and *C. baronessa*) may be attributed to benefits of assisted defense of dietary resources, and whether pair bond endurance enhances the effectiveness of assisted resource defense. Specifically, we aimed to test ARDH predictions that either: i) males primarily defend a feeding territory, and females benefit from male assistance by improved investment in feeding and energy reserves, or ii) both partners mutually defend their feeding territory and benefit from each other’s assistance by improved investment in feeding and energy reserves. If so, then finally, we tested the prediction that iii) pair bond endurance reduces intra-pair conflict and/or promotes assisted territory defense and/or energy reserves.

## Methods

### Study location and model species

This study was conducted on snorkel at adjacent sheltered reefs of Lizard Island, located on the northern Great Barrier Reef, Australia (14°40’S, 145°27’E) from January – March, 2014 at haphazard times between 0830-1730 hrs. Sampling was conducted on the two most locally abundant coral-feeding and pair bonding butterflyfishes, *C. lunulatus* and *C. baronessa* (Fig. 1). Only individuals that were within 80 % of the asymptotic size for the species (*C. lunulatus*: > 64 mm standard length (SL); *C. baronessa*: > 61 mm SL), and therefore likely to be reproductively mature^15^ were considered. Both species are territorial^39^ and are predominantly found in long-term, heterosexual partnerships^42,43^This study was conducted using Great Barrier Reef Marine Park Authority permits: G10/33239.1, G13/35909.1, G14/37213.1; James Cook University General Fisheries permit: 170251. It was approved by James Cook University Animal Ethics committee (approval # A1874), and performed in accordance with relevant guidelines and regulations.

**Figure 1.**
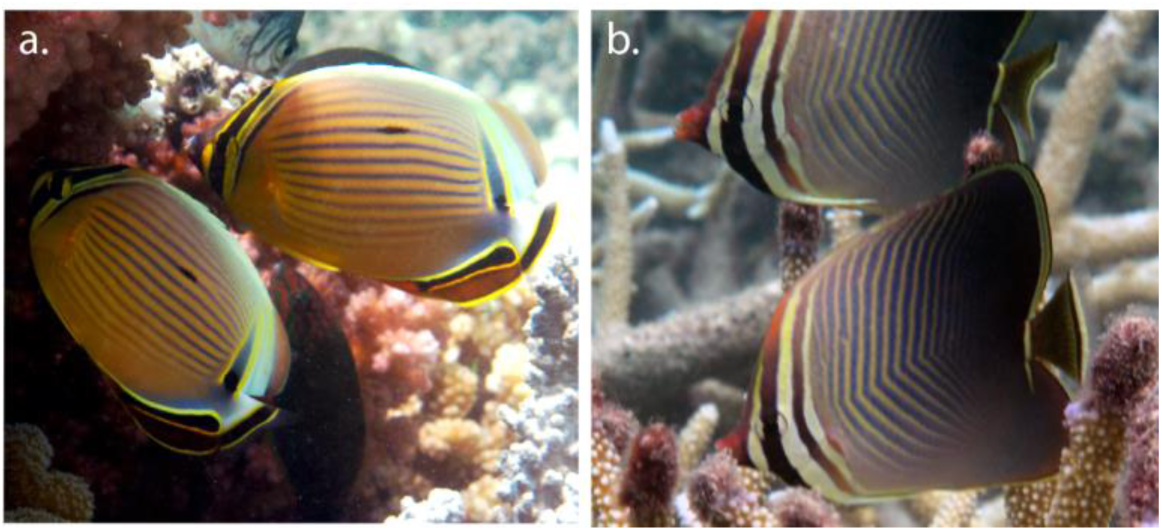
*Chaetodon lunulatus* (a) and *C. baronessa* (b) as models of pair bonding butterflyfishes used in the current study. At the study location, Lizard Island (GBR), these species are territorial coral feeding specialists that display enduring pair bonds. Pictures are of focal pairs, taken by J.P.N.

### Coordination and competitor aggression between male and female partners

To test whether pairs displayed either male-prioritized or mutual territory defence, we conducted *in situ* observations on naturally occurring paired and solitary individuals. Fishes were haphazardly encountered, approached from 2-4 m, and given 3-min to acclimate to observer presence. Following, social system was estimated during a 5-min observation. Pair bonded individuals were identified as displaying coordinated swimming exclusively with another conspecific, whereas solitary individuals were identified as displaying no coordinated swimming with another conspecific. For pair bonds, each individual was then identified using unique body markings (as per^31^) and assigned an identity number used for ongoing behavioral observations and sexing. Following, levels of coordination and competitor aggression were measured during a 6-min observation. Level of coordination was measured in pair bonded and solitary individuals, and determined by recording the presence or absence of coordinated swimming every 10-sec. Coordination, defined as the synchronisation of individuals’ movements in space and time^44^, was considered as the focal fish being positioned within a 2-m distance from another conspecific whilst being faced within a 315-45° angle relative to the faced position another conspecific (designated as 0°) (Fig. 2). Coordination was visually estimated after practicing accuracy on dummy fishes prior to the study. Level of aggression toward competitors was measured in male and female partners of pairs, and determined by quantifying the total number of aggressive acts (i.e., staring, chasing, fleeing, encircling, and head down, tail-up displays) expressed (see^45^ for detailed descriptions). Only aggression towards other butterflyfishes was measured, because for most butterflyfish, territorial competition is intra-familial^46^. While we attempted to measure aggression in both partners within a pair, there were few cases in which this occurred for only one partner. After each observation, both partners of pairs were collected by spearing through the dorsal musculature and sacrificed in an ice slurry for sex determination. A one-way ANOVA was used to compare the level of coordination within pairs between *C. lunulatus* and *C. baronessa*. Coordination data was square-root transformed prior to analysis to improve normality of residual variance. For each species, a one-way ANOVA was used to compare rate of aggressive acts between males and females within pairs.

**Figure 2.**
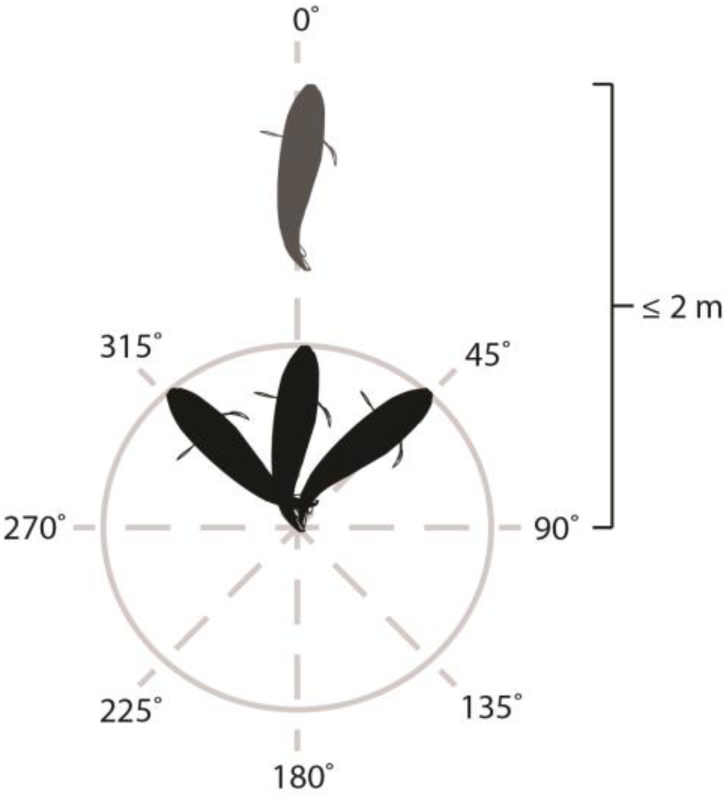
Coordinated swimming examined in pair bonded and solitary butterflyfish. Coordinated swimming by focal fish (black) was defined as being positioned within a 2-m distance from another conspecific (grey) whilst being faced within a 315-45° angle relative to the faced position of another conspecific (designated as 0°).

## Paired vs. solitary individuals

### Competitor aggression and feeding bites

To test whether one or both sexes benefit from pairing through reduced competitor aggression or increased feeding rates, we measured and compared these variables between naturally occurring paired and solitary individuals of both sexes. Individuals were considered pair bonded and solitary using the criteria previously described. After establishing their social status and undergoing 3-min acclimation to observer presence, focal individuals underwent a single 6-min observation to record: i) total feeding bite rate, determined by the number of bites taken on any coral ii) total feeding bites on preferred coral types (data on preferred coral food only collected for *C. baronessa*, which is *Acropora hyacinthus, A. florida,* and *Pocillopora damicornis*^47,48^), and iii) total rates of aggression toward neighboring butterflyfish. Rates of aggression may be affected by the local abundance of competitors (independent of levels of aggression exhibited by focal individuals), which was higher in paired than solitary fish. Therefore, in order to account for this potential confound, the number of aggressive acts recorded during replicate observations was standardized to per butterflyfish present within the immediate vicinity (3 m) of the fish’s feeding territory. Immediately following, butterflyfishes were collected by spearing through the dorsal musculature and sacrificed in an ice slurry for sex determination and energy reserves analysis. For each sex of each species, rates of aggression were compared between social conditions using non-parametric Mann-Whitney *U* tests, due to non-normal distribution of residual variance. For each species, feeding bite rate on total coral was compared between social conditions using a factorial ANOVA (with sex and social condition as fixed factors). For each sex of *C. baronessa,* a non-parametric Mann-Whitney *U* test was used to compare feeding bite rate on preferred coral between social conditions, due to non-normal distribution of residual variance.

## Enduring vs. new pairs: Intra-pair relations, and per capita competitor aggression and feeding bites

We used a partner removal-replacement experiment to examine whether pair bond endurance reduces territory defense or increases feeding of paired individuals by promoting cooperative territory defense and/or reducing intra-pair conflict. Both partners of naturally occurring pair bonds of *C. lunulatus* (n = 9) and *C. baronessa* (n = 10) were identified and monitored through time using unique body markings (as per^31^), which were photographed and printed on water-proof paper to assist observers (see Supplementary Figure S1 online for example photographs). Pairs were assumed to have been enduring, based on previous research showing a high level of partner endurance in these species at the study location^49^. Prior to experimentation, one individual from each pair was haphazardly chosen as the focal individual for the experiment. To identify the focal individual and its partners throughout the experiment, a photograph of both lateral sides of their body was taken, from which a unique body markings were recognized and used^43^. Behavioral expression of the focal individual while with its original partner was measured throughout an 8-min observation, for 5 consecutive days. Prior to each observation, the focal individual and its partner were allowed to acclimate to observer presence for 3-mins (as described above). During each observation, i) time spent coordinated swimming with partner, ii) aggression towards partner, iii) aggression per competitor, and iv) feeding bites of the focal individual were recorded using the methods previously described. Immediately following observations conducted over 5 consecutive days, the partner of the focal individual was removed via spearing and sacrificed in an ice slurry for sex determination and energy reserve analysis. All focal individuals had re-paired with a new partner within 18 hours of partner removal, as determined by identification methods previously described. We then conducted the same behavioral observations for a further 7 (*C. lunulatus*) or 9 (*C. baronessa*) consecutive days. After experimentation, the focal individual and its new partner were collected by spearing through the dorsal musculature and sacrificed in an ice slurry to determine the sex of both individuals and energy reserve of the focal individual’s new partner. Temporal changes in time spent coordinated swimming with partner, aggression towards partner, aggression per competitor, and feeding bites were analyzed using multivariate analysis of variance (MANOVA), with results displayed using canonical discriminant analysis (CDA)^50,51^.

## Solitary vs. newly paired vs. enduringly paired individuals: Differences in liver hepatocyte vacuolation

To assess changes in energy reserves in association with pairing and partner endurance, we compared liver hepatocyte vacuole density between individuals who i) naturally occurred in solitude (from *in situ* observation study), ii) were in new pair bonds (*C. lunulatus*: 7 day old partnerships; *C. baronessa*: 9 day old partnerships), and iii) were in naturally occurring enduring pair bonds (latter two conditions were acquired from individuals from partner removal experiment). Whole livers were dissected and fixed in 4% phosphate-buffered formalin (PBF). Fixed liver tissues were then dehydrated in a graded ethanol series and embedded in paraffin wax blocks. Tissues were sectioned at 5 µm, mounted onto glass slides, and stained using Mayer’s hematoxylin and eosin to emphasize hepatocyte vacuoles. Hepatocyte vacuole density was quantified using a Weibel eyepiece to record the proportion of points (out of 121) that intersected with hepatocyte vacuoles when viewed at X 40 magnification. Three estimates of hepatocyte vacuolation were taken for each of 3 cross sections, totaling 9 replicate estimates per fish liver, following^52^. In both species, differences in the percentage of liver hepatocyte vacuolation between solitary, newly paired, and enduringly paired fish were analyzed using a non-parametric Kruskal-Wallis one-way ANOVA^53^, due to non-normality in residual variance. Variation in hepatocyte vacuolation could not be analyzed for each sex separately, due to small sample sizes. Tukey and Kramer (Nemenyi) *post hoc* tests were used to identify differences between social condition means.

### Sex determination

The sex of focal fish was determined histologically. Gonads were removed and fixed in formaldehyde-acetic acid-calcium chloride (FACC) for at least 1 week. Thereafter, gonads were dehydrated in a graded alcohol series, cleared in xylene, embedded in paraplast, sectioned transversely (7 µm thick), and stained with hematoxylin and eosin. Sections were examined under a compound microscope (400 X magnification) for the presence of sperm (male) or oocytes (female)^15^.

## Results

### Coordination and competitor aggression between male and female partners

The two study species exhibited contrasting modes of cooperative or assisted territory defense. In both species, solitary individuals displayed no level of coordinated swimming with another conspecific. Pairs of *C. lunulatus* spent most of their time (56%) swimming with coordination throughout their feeding territory (Fig. 3a; see Supplementary Video S1 online for example). When encountering neighboring butterflyfishes, both partners displayed equal levels of aggressive acts (*F*_1,12_=1.01, *p* = 0.334; Fig. 3b), suggesting that there is mutual assistance in territory defense. By contrast, *C. baronessa* partners spent notably less time (10%) swimming with coordination (*F*_1,25_=40.04, *p* = 0.000) (Fig. 3a). In general, males tended to move over large distances within and along the boundaries of their territory whilst continuously foraging, whereas females tended to restrict movement to areas of their preferred coral (if available) whilst continuously foraging on the outcrop. When territorial disputes occurred, *C. baronessa* males exerted 42% higher levels of aggression than females (*F*_1,8_=7.51, *p* = 0.025; Fig. 3c), suggesting that in this species, territory defense is male-prioritized.

**Figure 3.**
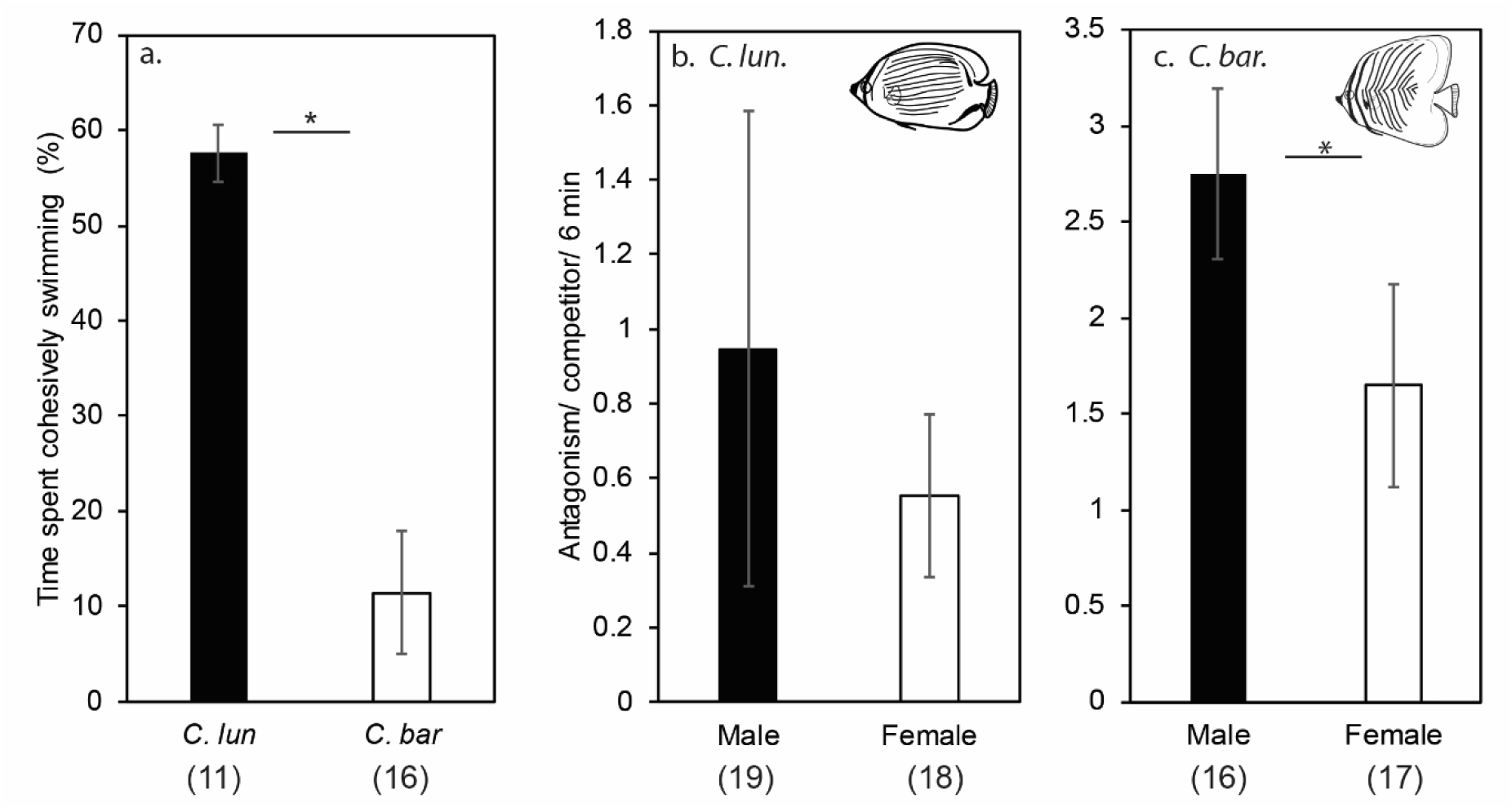
Patterns of (a) pair coordination and aggression towards competitors between male and female partners of (b) *C. lunulatus* and (c) *C. baronessa.* Data are represented as the mean ± SE; asterisks indicate statistically significant differences between treatment groups (ANOVA, *p* < 0.05). Sample sizes are listed below each treatment group.

### Paired vs. solitary individuals: Differences in competitor aggression and feeding bites

Across study sites, naturally occurring pairs of both species were common, whereas singletons were rare. A higher abundance of neighboring butterflyfish surrounding paired individual’s territories (in *C. lunulatus* by ~36%, in *C. baronessa* by ~75%) was found, suggesting that they had more neighboring competitors than solitary counterparts. For each sex of each species, after standardizing aggression to per competitor present, there was no apparent difference in rates of aggression between paired and solitary individuals (*C. lunulatus* males: z=-0.78, *p*=0.48; females: z = -1.69, *p*=0.11; *C. baronessa* males: z=-0.53, *p*=0.64; females: z=-0.00, *p*=1.00; Fig. 4a, c).

**Figure 4.**
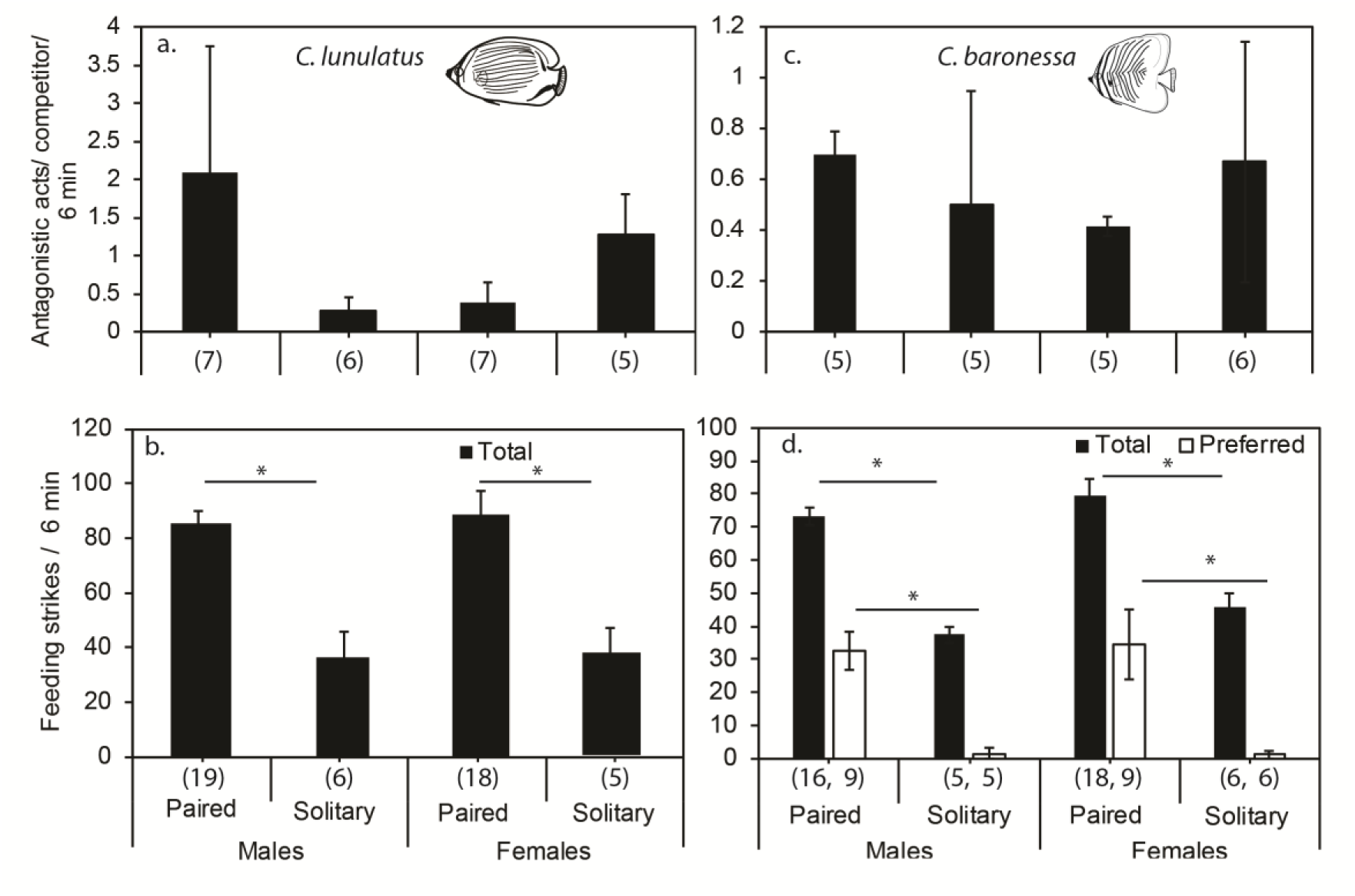
Differences in (a, c) aggression towards competitors and (b, d) bite rates between paired and solitary *C. lunulatus* and *C. baronessa* individuals. Total feeding strikes refer to the number of bites taken on any coral, whereas preferred feeding strikes (only measured in *C. baronessa*) refer to number of feeding bites taken on preferred coral types (i.e., ***Acropora hyacinthus, A. florida,* and *Pocillopora damicornis***). Data are represented as the mean ± SE; asterisks indicate statistically significant differences between treatment groups (ANOVA or Mann-Whitney *U*, *p* < 0.05). Sample sizes are listed below each treatment group.

In both sexes of both species, paired individuals have higher feeding bite rates than solitary counterparts (*C. lunulatus*: total coral bites: *F*_1,44_= 28.57*, p* = 0.00; *C. baronessa*: total coral bites: *F*_1,40_= 28.91*, p* = 0.00, preferred coral bites: z=-2.42, *p*=0.013 (males), z=-2.88, *p*=0.00 (females), Fig. 4b, d). In *C. lunulatus,* single males took 36.83 ± 8.87 SE bites per 6-min, whereas paired males took 86.21 ± 4.28 SE bites (approx. 57 % more); and single females took 37.40 ± 9.41 SE bites per 6-min, whereas paired females took 88.67 ± 8.56 SE bites (approx. 58 % more). Consistently, in *C. baronessa,* single females took 46.00 ± 3.21 SE total coral bites per 6-min, among which 1.17 ± 1.17 SE bites were on preferred coral; whereas paired females took 79.67 ± 4.91 SE total coral bites per 6-min, among which 33.33 ± 10.98 SE bites were on preferred coral (~43 % more total coral bites, and ~96% more preferred coral bites). Similarly, single males took 37.40 ± 3.93 SE bites per 6-min (1.4 ± 1.16 SE bites on preferred coral), whereas paired males took 73.13 ± 4.86 SE bites per 6-min (32.44 ± 10.54 SE bites on preferred coral), equating to ~49 % more bites on coral and ~96 % more bites on preferred corals.

### Enduring vs. new pairs: Intra-pair relations, and per capita competitor aggression and feeding bites

#### Costs of new pairing

Throughout the 5 consecutive days leading up to partner removal, all focal individuals maintained their same partner and territory. Within 18 hours of removing their original partner, all focal individuals had kept their same territory, where they had re-paired with a new partner. After re-pairing, the activity profile (union of pair swimming, within-pair aggression rate, competitor aggression rate, and feeding bite rate) of fishes dramatically changed (*C. lunulatus:* Pillai’s trace = 0.23, df = 1, *p* < 0.001; *C. baronessa*: Pillai's trace = 0.29, df = 1, *p* < 0.001; Fig 4a, b canonical score plots). When individuals formed new partnerships, they mostly displayed higher within-pair aggression, and to a lesser extent altered coordinated swimming (*C. lunulatus*: lower coordinated swimming; *C. baronessa*: higher coordinated swimming), higher aggression per neighbouring competitor, and lower feeding bites than when they were in their enduring partnership (Fig. 5a, b canonical structure plots; see Supplementary Video S2 online for example of day 1 of new partnership).

**Figure 5.**
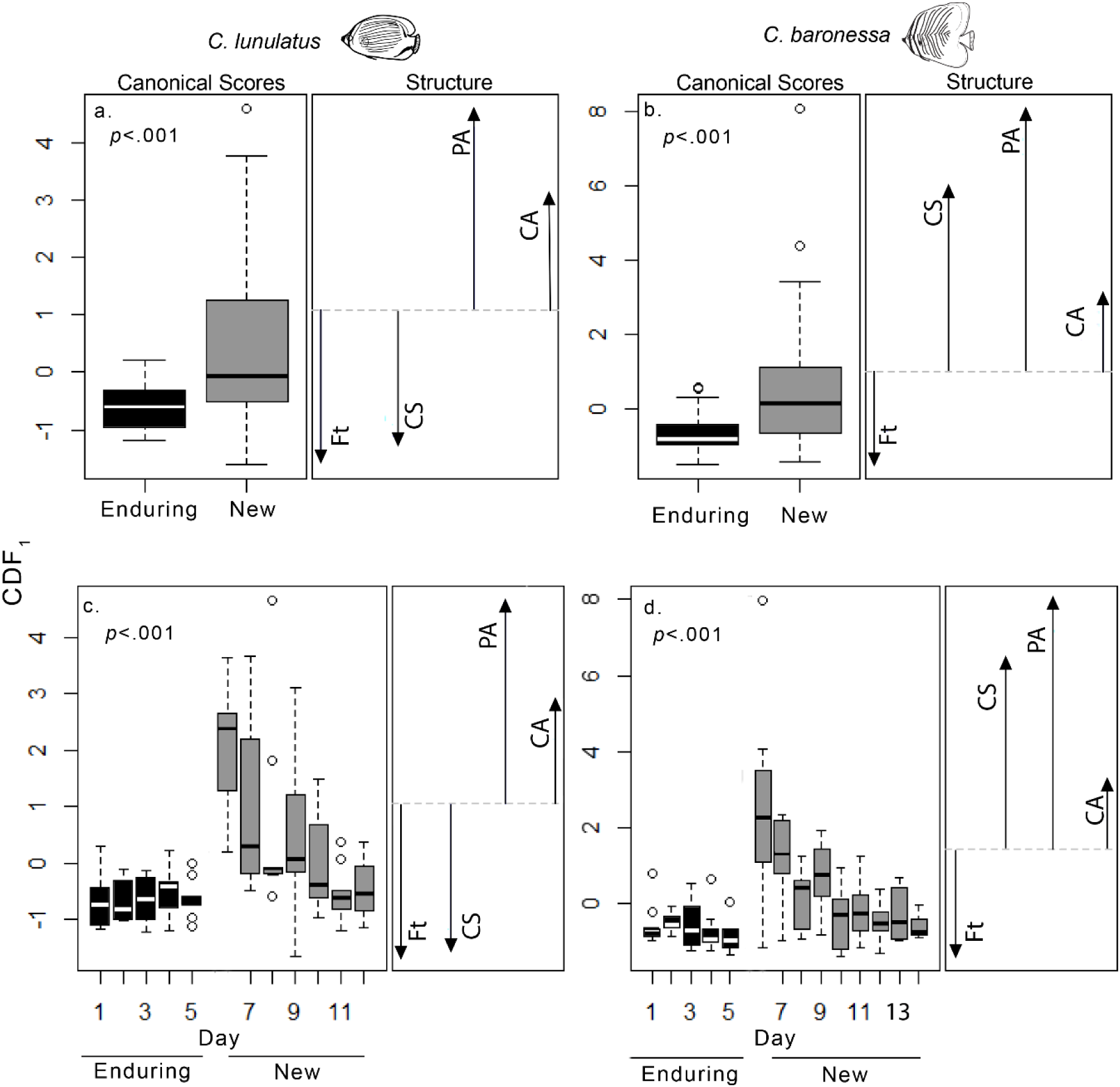
Changes in intra-pair relations, aggression towards competitors, and feeding bites in response to (a, b) re-pairing, and (c, d) subsequent endurance of new pairs throughout several days. Means of standardized canonical scores of the first canonical discriminant function (CDF_1_) are represented by box and whisker plots. Structure vectors show the relative strength (length of the vector relative to length of other vectors) and direction (+ or -) of the correlation between each contributing response variable and the canonical discriminant function. MANOVA p-value for change in activity profile in response to relationship phase or day is shown in the corner. (a,b) In both species, re-pairing with a new partner increases intra-pair aggression (PA). Concurrently, it (a) reduces coordinated swimming (CS) in *C. lunulatus* (n = 9), and (b) increases coordinated swimming in *C. baronessa* (n=10). (a,b) These changes in intra-pair relations are associated with increased competitor aggression (CA) and a reduction in total feeding bites (Ft). (c,d) However, as new pairs endure, intra-pair relations recover along with recovered losses in competitor aggression and feeding bite efficiency.

#### Recovery with new partnership endurance

After re-pairing, focal individuals maintained association with their new partner throughout the remainder of the study (*C. lunulatus,* seven days; *C. baronessa,* nine days), except for one *C. lunulatus* individual, who underwent a second re-pairing 3 days after its original partner was removed. As new pairs endured, focal individuals’ activity profiles significantly changed (*C. lunulatus*: Pillai's trace = 0.22, df = 1, *p* < 0.001; *C. baronessa*: Pillai's trace = 0.23, df = 1, *p* < 0.001; Fig. 5c, d canonical score plots). This change was mostly attributed to a reduction in intra-pair aggression, and to a lesser extent to adjusted coordinated swimming (*C. lunulatus*: increased; *C. baronessa*: decreased), reduced aggression per competitor, and increased feeding bites (Fig. 5c, d canonical structure plots). Notably, after 4 days of new pair endurance, these behaviors steadily recovered to levels displayed by original pairs (Supplementary Tables S3 and S4; Fig. 5c, d canonical structure plots).

### Energy reserves of solitary vs. newly paired vs. enduringly paired individuals

For both *C. baronessa* and *C. lunulatus*, liver hepatocyte vacuole density varied significantly with social condition (*C. lunulatus*: Kruskal-Wallis = 19.39, df = 2, *p* < 0.001; *C. baronessa*: Kruskal-Wallis = 10.27, df = 2, *p* = 0.006). While there was no difference in liver vacuole density between individuals that were in enduring and relatively new (i.e., that persisted for 7-9 days) partnerships, paired individuals had greater hepatocyte vacuolation than solitary counterparts (Fig. 6a, b).

**Figure 6.**
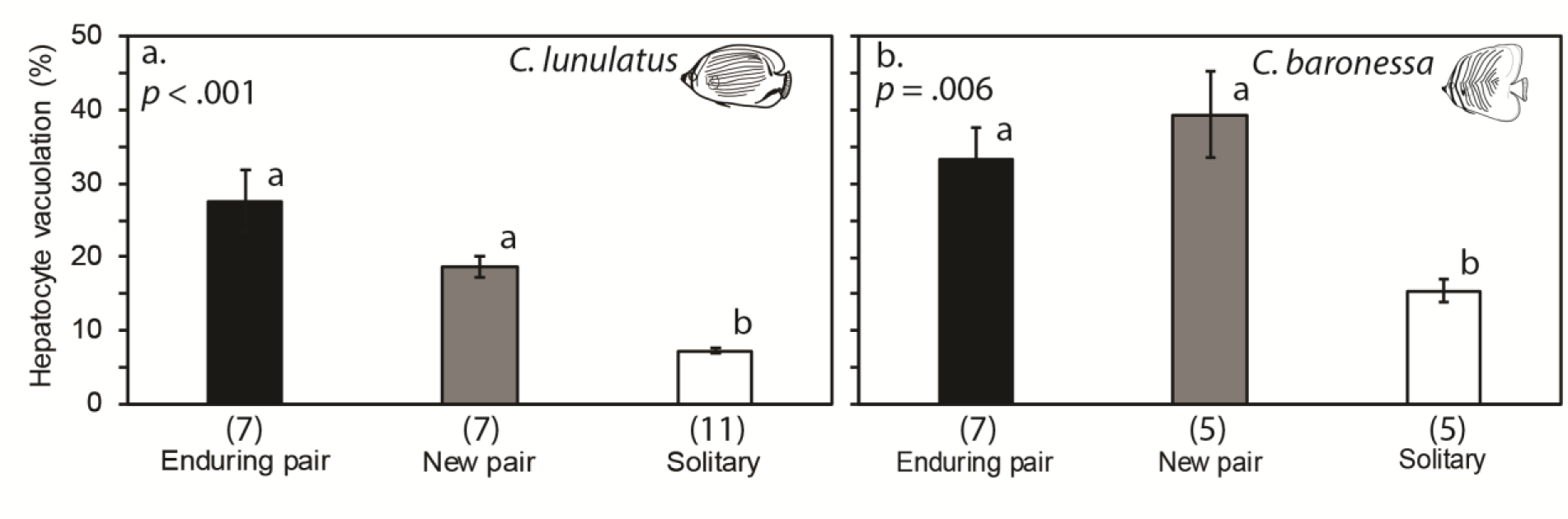
Variation in liver hepatocyte vacuole density among individuals in enduring partnerships, new partnerships (7-9-day persistence), and living in solitude. (a) *C. lunulatus;* (b) *C. baronessa.* **Figure 5 text:** Data are represented as the mean ± SE. Kruskal-Wallis *p* value is shown in top corners, while groups not sharing the same letter are significantly different [Tukey and Kramer (Nemenyi) post-hoc analysis at *p*< 0.05]. Sample sizes are listed below each treatment group.

## Discussion

In this study, we provide field-based observational evidence for the ARDH for pair bonding in two species of butterflyfishes. We show that pairs of *C. baronessa* and *C. lunulatus* appear to exhibit alternative modes of assisted territory defense (male-prioritized and mutual defense, respectively) that is associated with increased feeding bites relative to solitary counterparts. We furthermore provide the first evidence that this feeding bite advantage translates into significant gains in energy reserves in butterflyfishes. Finally, this is one of the first studies to examine possible reason(s) for partner endurance within the context of ARD in animals, providing experimental evidence that partner endurance plays a critical role by inhibiting conflict and promoting ARD between partners.

It has previously been proposed that, where ARD drives pairing, males nonetheless take-on the greatest burden of defense^54^. Although this is largely consistent with our results for *C. baronessa*, our findings for *C. lunulatus* contribute to a growing body of literature indicating that males and females may contribute equally to resource defense, because both may equally benefit from each other’s assistance (Supplementary Table S1 online). Partners frequently swam with coordination while foraging throughout their territory. When encountering neighboring butterflyfishes, aggression was generally passive, consistent with some other butterflyfishes^38,41^. Presumably, the function of pair swimming in butterflyfish pair bonds may be akin to duetting in bird pair bonds, in that it conspicuously advertises territory occupancy, thereby avoiding usurpation attempts by neighbors^11^. Notably, when territorial aggression did occur, it was exerted by both partners mutually. For both sexes, this ostensive co-defense in paired individuals was associated with an improved feeding bite rate (by ~ 58 %), and energy reserves (by 69 %, as indicated by hepatocyte vacuolation) relative to solitary counterparts. Consistently, in *C. chrysurus* (= *paucifasciatus*), male-female partners continuously travel closely together throughout their territory, mutually engage in territory defense, and both partners have higher feeding rates than solitary individuals^11^.

In contrast to *C. lunulatus*, territory defense by pairs of *C. baronessa* appeared to be male-prioritized. Partners frequently traveled independently from each other, spending only ~10 % of their time swimming with coordination. For most time (~90 %) males independently patrolled larger areas within and along the boundaries of territories, exerting ~ 42 % more aggression towards neighboring butterflyfishes than females. This seemed to allow females to mainly focus on foraging, notably in a more restricted area within the territory that contained a dominant assemblage of preferred coral (e.g., *A. hyacinthus* or *A. florida*). In association, paired females bit 43% more total coral, and 96% more preferred coral then their solitary counterparts. Moreover, and probably because of increased food bites, liver lipid reserves in paired individuals were higher by ~ 57 % than solitary individuals, though no distinction was made between males versus females. Among species that pair for assisted resource defense purposes, male-prioritized defense is the most commonly reported mode of assistance (Supplementary Table S2 online), where it has been attributed to supporting female food consumption in birds^55,56^ and butterflyfishes^32^, presumably enabling the male to share his mate’s subsequent increased fecundity^32,56^. This form of sexual division-of-labor is thought to occur when female egg production is especially costly, and disproportionally more male assistance is required for females to build the energy reserves needed for egg production^32,57^. Egg production in female *C. baronessa* may be particularly energetically costly, thereby favoring male-prioritized over mutual defense. Although some of *C. baronessa*’s preferred diet, *Acropora* corals, provide the best energetic return among coral families (after accounting for feeding efficiency)^58,59^, they nonetheless exhibit higher feeding rates on coral tissue per day than *C. lunulatus* and other congeners^59^, indicating that perhaps they have a relatively low energetic absorption efficiency. To ascertain this possibility, analysis of energetic absorption efficiency (relative to other corallivorous species) would be required. Interestingly, paired *C. baronessa* males bit 49 % more total, and 96 % more preferred coral than their solitary counterparts. This might suggest that in addition to presumably sharing in female’s increased fecundity, pairing males might also directly confer an advantage to food consumption, due to the (albeit relatively little) territorial defense assistance provided by females.

In previous studies of pair bonding butterflyfishes, ARD led to increased feeding rates by reducing the time needed for territory defense, thereby providing more time to invest in feeding^11,40^. In this study, however, we found no evidence that paired individuals could effectively defend territories with lower levels of aggression relative to solitary counterparts. Importantly, rates of aggression tended to be highly variable. This, coupled with the relatively low sample sizes, limited power to detect potential differences in aggression between paired and solitary individuals. Alternatively, ARD may have enabled pairs to establish territories with greater food availability, thereby providing feeding and energy reserve benefits independently of reduced per capita territory defense. It may also be argued that rather than ARD, increased feeding and energy reserves might be consequent of other function(s) of pairing (e.g., increased predator vigilance)^15,60^. Finally, it is conceivable that higher food consumption and energy reserves promoted pair bonding. Nevertheless, a causal effect of ARD on improved feeding experimentally shown in other pair bonding butterflyfish species^11,61^ further supports the idea that it also exist in species of the current study. To be certain, however, this should now be confirmed experimentally.

Although species who pair for ARD display long-term partner fidelity, reasons for this are almost wholly unknown (Supplementary Table S2 online). In the current study, we show that within 18 hours of removing their original partner, all remaining fish had kept their same territory, wherein they had re-paired with a new partner. This indicates strong territory fidelity. Moreover, while there is strong pressure to be paired, this is not due to partner/mate scarcity. There are, however, definite benefits of pair bond endurance. Experimentally inducing new partnerships caused an immediate and marked decline in partner relations, although these were relatively short-lived. This was primarily driven by increased in intra-pair conflict, and to a lesser extent by reduced expression of species-specific modes of ARD, as indicated by decreased pair swimming in *C. lunulatus*, and increased pair swimming in *C. baronessa*. This decline in partner relations appeared to initiate heightened territorial activity with neighbouring butterflyfish pairs. Subsequently, newly paired individuals suffered from having to shift investment from feeding to territory defence aggression. As new partnerships subsequently endured, however, these intra-and inter-pair disruptions abated, and incurred costs to individual territory defence-feeding budgets recovered accordingly. Similarly, it has been shown for wood louse (*Hemilepistus reaumuri*) that widowed individuals with established territories will initially aggressively resist the elicitation to form new partnerships prior to conceding^19^. It has also been shown that pair bonds of longer duration monopolize higher quality feeding territories, ostensibly through enhanced co-operation, and this is further linked to improvements in life-time reproductive success (barnacle geese, *Branta leucopsis^22,56^*). Overall, our results suggest that partner fidelity is exhibited in *Chaetodon* butterflyfishes because it plays a critical role in promoting assisted resource defence, and inhibits intra-and inter-pair conflict, ultimately conferring feeding investment gains. Although we found no evidence that this translates into energy reserve gains, this may be consequent of limited sample size and/or sampling after new pairs had already endured for 7-9 days, when they displayed fully-recovered behavioral and energetic profiles. Indeed, in fishes, liver hepatocyte vacuole density has been shown to respond rapidly to changes in feeding (i.e., within 2 days)^62,63^. To ascertain this, energy reserves should be re-sampled in more individuals and on a shorter time-scale throughout the development of new partnerships. Of course, there may be other reasons for long-term partner fidelity among pair bonding species who exhibit ARD. These might include partners experiencing a delay in the time at which their services are reciprocated^3^. For example, if male assistance is based on increasing female feeding to share her improved fecundity, then males may remain with females across reproductive periods if there is a time-lag between enhanced female feeding and egg production^3^. Partner fidelity may also be attributed mutual site-attachment to the feeding territory, which may arise if it new territories are scarce or competitively costly to acquire^18,38^.

How and why might partner fidelity promote assisted resource defense and inhibit intra-pair conflict in these species? Perhaps partner fidelity improves assisted territory defense through partner familiarity. Indeed, it has been shown in fishes that cooperation with specific partners stabilizes over time, because individuals are more cooperative with familiar partners^64^. The mechanism(s) for this may be unique to the species-specific mode of cooperative assistance. For *C. lunulatus*, partners appear to work together simultaneously to provide mutual assistance, and as such, familiarity may facilitate learning and accurate prediction of partner behavior (e.g., chosen defense route or routine), thereby fine-tuning pair-wise coordination^65–68^. For *C. baronessa*, partners appear to exhibit male-prioritized assistance in exchange for sequentially reciprocated partitioning of services/resources by females (i.e., direct reciprocity), and as such, partner familiarity may allow individuals to learn which “partner control mechanism” is best suited to stabilize cooperation, based on the tendency of partners to reciprocate (or cheat) in the past^69,70^. Upon new pair formation, partner familiarity (and therefore effective co-operation, and co-operatively derived feeding benefits) takes several days to develop; however, the cost of food sharing is immediately incurred. Hence, until co-operative relations develop, the costs (food sharing) likely outweigh the benefits (maximizing feeding investment) of pairing, causing territory holders to aggressively resist new partner elicitation.

Energy acquisition is fundamental to growth, reproduction, and maintenance for all animals^71^. However, corallivorous butterflyfishes rely almost exclusively on a diet of hard coral^72^, which is a relatively nutrient poor, but abundant resource^33,37^. Consequently, both sexes are energy maximizers, feeding almost continuously^32,38,40^. Foraging is therefore constrained by time spent on other activities, including territory defense. As such, attributes that alleviate time constraints on foraging are likely to directly benefit individual fitness^32^. The study suggests that partners of *C. lunulatus* and *C. baronessa* display territorial defense assistance to increase their feeding of coral food and energy reserves. We further show that partner fidelity plays a critical role in this function by inhibiting intra-and inter-pair conflict and promoting territorial defense assistance between partners, providing an ecological advantage to pair formation and fidelity in these species. Whether this translates into an adaptive advantage should now be addressed by undertaking long-term monitoring studies to discern whether long-term pair bonding also confers relatively higher survivorship and/or life-time fitness benefits^56^.

## Acknowledgements

We thank Manuela Giammusso, Patrick Smallhorn-West, Andrés Jácome, and staff at Lizard Island Research Station for field assistance. We thank the anonymous reviewer(s) for providing helpful feedback on the manuscript. We acknowledge with gratitude the fishes that were used in the current study. This research was supported by an ARC research grant to M.S.P and S.P.W.

## Author contributions

J.P.N, S.P.W and M.S.P designed the study; J.P.N., D.J.C, and K.J.N. collected the data; J.P.N., S.P.W. and K.J.N. analyzed the data, J.P.N. wrote the manuscript, and all authors contributed to reviewing and editing the manuscript.

## Additional information

### Supplementary information

See accompanying documents.

